# Western Washington culvert prioritization indexes: A cross jurisdictional comparative analysis of management strategies for salmon recovery

**DOI:** 10.1101/2022.11.07.515553

**Authors:** Catalina A. Burch, Sunny L. Jardine, Connor Lewis-Smith, Braeden Van Deynze

**Author notes:** Corresponding author. Catalina Burch. Washington Department of Fish and Wildlife (Braeden Van Deynze).

## Abstract

Culverts throughout western Washington state, U.S.A., contribute to declines in anadromous fish populations by blocking fish passage. A federal injunction requires that Washington state must restore 90 percent of blocked habitat caused by state-owned culverts by 2030. This ruling prompted the development of numerous prioritization indexes (PI), the ranking of barriers through weighted combinations of barrier-specific metrics, by culvert owning entities (i.e., counties, cities, ect.) within the injunction area. We conduct a comparative analysis of PIs within the injunction case area, investigating their ability to distinguish between barriers, their transferability in terms of scoring metrics utilized, how scoring weights differ, and the preferences implied thereby. We document the use of six distinct PI methods by ten barrier owning entities, and find that some PIs used many shared metrics, while others used a high percentage of unique metrics that would be difficult to replicate outside the entity’s jurisdiction. While habitat quality, habitat quantity, and connectivity were considered across all PIs, we found a high level of variation in terms of the metric weights. Our methods can be employed in other geographies or for other restoration PI planning efforts, and our results may facilitate the development and refinement of future PIs.

## Introduction

Across the United States there are millions of culverts at road-stream crossings, allowing for safe transport across waterways and flood management (Martin 2019, McKay et al 2020).

However, road culverts often create barriers for fish passage and can fragment habitat (Januchowski-Hartley et al 2013), which is an issue of particular concern for migratory species such as salmonids (Nathan et al 2018). For example, barrier culvert habitat fragmentation has been found to limit salmon access to critical spawning and rearing habitat and reduce genetic exchange over time (Sheer & Steel 2006; Tudorache et al 2008).

In Western Washington state, fish passage barrier correction projects are accelerating as state and local governments respond to a unique political and policy landscape (Blum 2017, Roni et al 2002). In 2007, a group of 21 indigenous tribes began settlement negotiations with Washington state concerning barrier culverts that infringe upon their treaty rights to take fish in their usual and accustomed areas (Blum 2017). However, negotiations failed, which led to a permanent injunction issued by the federal court, in favor of the tribes, mandating that Washington Department of Fish and Wildlife (WDFW) and Washington Department of State Transportation (WSDOT) restore 90 percent of fish habitat blocked by outdated culverts by 2030 (*United States v. Washington* 2017). The injunction Case Area includes the lands and waters ceded by the Tribes to the state of Washington in exchange for guaranteed fishing rights in the Stevens Treaties (*United States v. Washington* 2017). While the injunction only applies to state-owned culverts, 16,185 additional barrier culverts owned by a variety of different entities (i.e., cities, counties, private landowners) have been identified within the Case Area, and many of these entities have also accelerated barrier correction with support from both local and state funding. This ruling created a unique political environment that prompted the rapid development of culvert inventories and prioritization strategies within the Case Area.

In Washington state and beyond, natural resource managers often develop and apply prioritization strategies to guide barrier culvert removal and replacement (McKay et al 2020). To develop a plan, managers must have a reliable inventory of culverts within their entity, which requires advanced mapping capabilities and extensive ground truthing efforts (Kemp & O’Hanley 2010, McKay et al 2017). Barriers in an inventory are then commonly assessed using quantitative and qualitative metrics, which typically consider habitat quantity and quality, connectivity and passability, cost, and species-specific metrics (Martin 2019), to identify high priority barriers for correction (Kemp & O’Hanley 2010, McKay et al 2017). Specific metrics used to quantify these criteria vary, and entities can have distinct priorities which are reflected in their chosen methods.

There are many different potential prioritization approaches including mathematical optimization, scoring and ranking, selecting projects based on expert knowledge, and reactively responding to opportunities presented through either auxiliary construction or funding (McKay et al 2020). Prioritization Indexes (PI), the most commonly employed method (Kemp & O’Hanley 2010), use a weighted ranking of multiple metrics to score individual culverts which are candidates for restoration (McKay et al 2020). The PI approach is advantageous because it is flexible to multiple objectives, does not require mathematical expertise, is not a “black box”, and facilitates stakeholder buy in (McKay et al 2020).

Noted limitations of PIs include their inefficiency if not considering connectivity between barriers and the interdependency of barrier removal efforts (McKay et al 2020). For example, restoring a high scoring upstream barrier will not lead to habitat gains unless all barriers downstream are also restored. Additionally, metric weighting is highly subjective and requires a lengthy recursive development process (McKay et al 2020). However, for many entities the advantages of PIs outweigh their limitations and as such are commonplace. Thus it is important to understand what metrics are used, how they are weighted, and the implicit trade-offs in different PI equations, particularly in Western Washington where barrier correction efforts are rapidly accelerating in response to the injunction.

There are several synthesis papers exploring fish passage prioritization methods in the literature (Kemp & O’Hanley 2010, McKay et al 2017, McKay et al 2020). In addition, there is ample literature describing the development process of designing a PI (Hoenke et al 2014, Mader & Maier 2008, Martin 2019), and comparing the PI approach to other prioritization strategies (Kemp & O’Hanley 2010, McKay et al 2020, Roni et al 2002). Previous comparative research has investigated prioritizations across large spatial scales: globally (Kemp & O’Hanley, McKay et al 2020, McKay et al 2017), nationally (NOAA 2021), and across multiple states (Martin et al 2019, Roni & Beechie 2002). Prioritization research on smaller scales (Hoenke et al 2014, Mader & Maier 2008), have focused on the development of an individual PI within a region, but not the comparison of multiple PIs. Additionally, the literature is lacking information on the preferences and tradeoffs implied by the construction of PIs using different methods.

Here we summarize and analyze barrier culvert PIs developed within the Case Area since the 2013 injunction. This study capitalizes on the rare opportunity to study multiple PIs created within in a well-defined timeframe and geographic region and addressing a single conservation issue. Comparing PI methods across entities improves transparency and understanding of the implications of various PI methods, while potentially facilitating the sharing of data and methods. This builds upon existing work to facilitate coordination and communication within barrier removal efforts (McKay et al. 2020). This research is especially timely in light of the culverts injunction and recent federal infrastructure funding, which accelerates ongoing barrier correction efforts (Cordan & Bradley 2021).

Our cross jurisdictional comparative analysis addresses the following questions: (1) How clearly do PI scores distinguish high scoring barriers from low scoring barriers? (2) How transferable are the PI scores in terms of scoring metrics utilized? (3) How do PI metric scoring weights differ? (4) What are the tradeoffs between objectives implied by the various PI equations? We identified six entities who developed unique PIs within the Case Area, and four additional entities that share a PI method developed by a state agency.

We found that PI scoring methods are publicly available though often buried within lengthy technical reports. This article addresses this issue by synthesizing across entities to provide detailed information on PI equation metric inputs and weighting. Among the PIs studied, we found a high level of variation in terms of goals, geographic area, culvert ownership type, and metrics employed. We found that all PIs, except one, have non-normal positively skewed distributions which clearly distinguish high priority barriers. Some PIs used more transferable methods with many shared metrics, while others used a high percentage of unique metrics that would be difficult to replicate outside the PI geography. One common theme is that habitat quality, habitat quantity, and connectivity are considered across all linear PIs. Among the PIs studied, we found a high level of variation in terms of the relative weight given to each metric. The methods we present are applicable to other geographies or for other restoration PI planning efforts, and our results may facilitate the development and refinement of future PIs.

## Methods

### Study Area and Systematic Review

We defined our study area as the Washington Culverts Injunction case area (Figure 1), hereafter referred to as the Case Area. Within this area, we conducted a directed internet search for documentation of culvert PIs published since the 2013 injunction. Specifically, a three-phase search-and-screen procedure, adapted from Prokopy et al. (2019), was employed to identify all uses of PI methods to score-and-rank barrier culverts.

**Figure 1.**
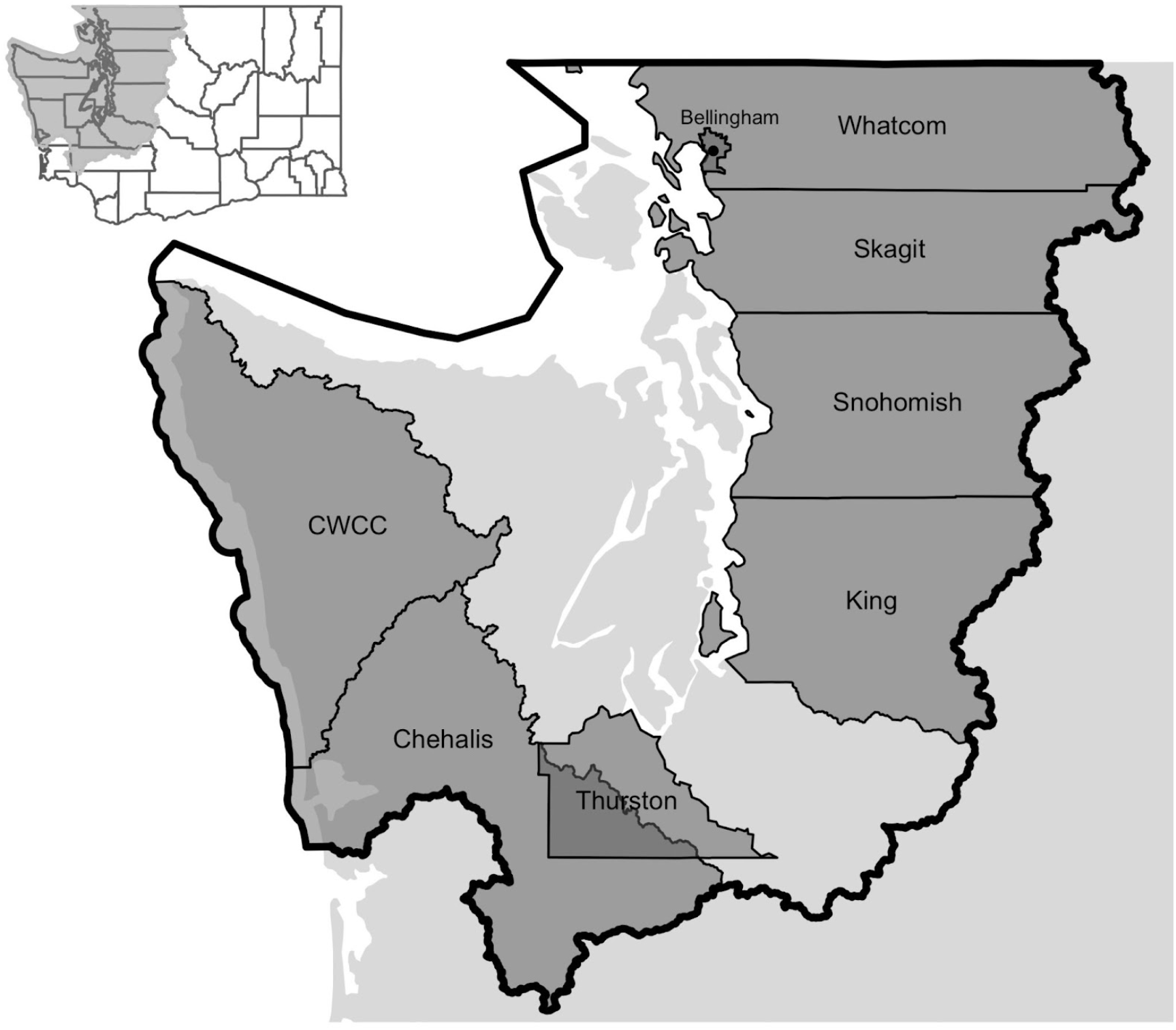
Map of study area showing the case area within the state of Washington. Each PI geography is shown within the case area. Shading was used to denote PIs that overlap geographically. Regions where no PI was located are shown as an unlabeled aggregated land mass. WDFW and WSDOT are not shown because their geographies span the entire case area.

The first screen was a multilayered internet search for specific keywords conducted through a general Google search on January 10, 2022, for the keywords “Washington culvert prioritization,” and selecting the first 20 search results to explore for documentation of an existing PI program. For each search result, we used the Find function to search for the inclusion of the word “prioritization.” Using this method, we identified eight entities (two state agencies, two non-profits, one basin effort, two counties, and one city) that matched the search criteria.

The next search was a county webpage search, completed for each of the 14 counties within the Case Area, on January 17, 2022. We searched for the keyword “culvert prioritization” on each county’s website and selected the first 10 search results to further explore, which uncovered one additional county entity that matched our search criteria.

The second phase involved reviewing the webpages for the ten entities utilizing or developing PIs to identify professional points of contact, supplementary reports, and data sources, and to obtain data describing the PI scores and contributing metrics when possible. Our team reached out via email to lead authors and referenced contact persons identified via jurisdictional reports or websites to acquire additional data, reports, and presentations for review.

The third phase involved review of materials, where two of our team members read the screened reports and presentations, followed by quality checks for the barrier data. Missing or incomplete information was noted.

### Data Assembly

We synthesized publicly available information and datasets provided by personal communications to produce a table with the following fields: PI developer (entity), ownership extent, geographic extent, prioritization type, number of barrier culverts with PI scores, and use of interactive tools (Table S1). A secondary detailed summary table includes equation type, metric inputs, and metric definitions (Table S2). The purpose of these summary tables was to compile important information and identify any gaps in our data.

### Analysis

#### All PIs: Distribution Analysis

We began with a distribution analysis for each PI to better understand the spread of scores generated by each method and how clearly high scoring barriers are distinguished from low scoring barriers. We expected that the ideal distribution would be non-normal with a positive skew making it easier to distinguish a clear hierarchy of priority barriers where correction efforts can begin. However, we also recognize that some entities may have chosen to produce a coarse hierarchy of barrier scores that are then further screened with methods outside of the PI score, e.g., through opportunistic or expert decision making (Pressey & Bottrill 2008).

We first summarized barrier PI scores separately for each jurisdictional entity (entity). Data from WDFW, the city of Bellingham, Chehalis Basin, Cold Water Connection Campaign (CWCC), Thurston County, and King County were filtered to only include barrier culverts (<100% fish passability) that received a PI score. Dams, fish screens, natural obstructions and other barriers were excluded from analysis.

We then calculated summary statistics on the PI scores of barrier culverts from each entity and conducted Shapiro Wilk tests for normality. To facilitate comparison across entities, PI scores for each entity were normalized to a scale of 0 to 100.

#### All PIs: Metric Categorization

We categorized all metrics contributing to PI scores for each entity to measure how implied priorities vary across the Case Area in terms of metrics considered. To compare metrics across entities we categorized them according to stated objectives (Table 1). These objectives were determined through the description of the metrics in prioritization reports and manuals, where possible, and through identifying qualitative similarities where documentation was missing.

**Table 1.** Definitions of metric categorizations.

| Category Name | Definition |
| --- | --- |
| Connectivity | Metrics that consider fish passability through the barrier, and the number/passability of upstream or downstream barriers. |
| Coordination | Metrics that include coordination with other projects, community support or impacts, and expert opinion. |
| Cost | Metrics that estimate the cost of the project and potential funding opportunities. |
| Future Conditions | Metrics that consider future habitat and environmental conditions. |
| Habitat quality | Metrics that include current suitability of habitat or environmental conditions at/upstream/downstream from the barrier. |
| Habitat Quantity | Metrics that include upstream habitat gain from barrier removal. |
| Species Metrics | Metrics that factor in species specific or life history characteristics. |

#### All PIs: Shared Metric Analysis

To assess transferability of scoring methods outside of each entity, we used a qualitative comparison matrix to identify which metrics were shared between two or more PIs. Metrics that were labeled shared measured the same type of attribute, such as number of downstream barriers or distance of upstream habitat, but may have used different methods or units for measuring that attribute. For example, one PI could have measured downstream barriers as a binary (yes/no) metric, while another PI measured the total quantity of downstream barriers. Metrics that were shared by at least one other PI were coded “shared,” and metrics that were not shared by any PIs within the Case Area were coded as “unique” metrics. We report the percentage of shared metrics for each PI in total and broken out by each metric category.

#### All PIs: Formula Categorization

We categorize the PI methods according to whether the scores combine metrics through linear (i.e., weighted sum) or non-linear (i.e., the weight of one metric depends on the value of another) combination. The distinction is important because metric weighting and implied tradeoffs cannot be directly compared across formula types. Our review identified both linear PI formulas and non-linear PI formulas. For the remainder of the analysis, we compare weights of the linear PIs before providing a separate analysis of the behavior of the non-linear PI method under different representative scenarios.

#### Linear PIs: Category weights

By sorting metrics for all PIs into categories we can directly compare the weight of different categories across all methods to understand how implied priorities vary. Metric weighting was identified for each PI through personal correspondence or technical reports. Linear PI scores are of the general form:

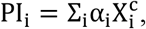

where the α_i_ are weights on individual metrics 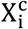, and superscript *c* denotes the category of metric *i* (see Table 1). However, many of the linear PI formulas include multiple metrics that contribute to the same objective. Thus, to quantify the weights that entities assign to objectives, we calculate the scoring weight of each category according to:

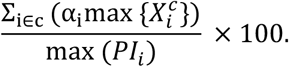

#### Non-Linear PIs: Weights

We identified one non-linear PI equation in the Case Area, developed by WDFW and used by four entities:

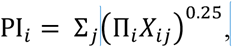

where the individual PI sums over each species the quadratic root of the product of species-specific metrics *X*_*ij*_.

Unlike linear formulas, the impact of increasing a metric by one unit changes based on the product of values of the metric and all other metrics in the formula, which makes it challenging to assess the weight that managers place on a single objective. Thus, to understand the implicit tradeoffs of the non-linear PI formula, we examine how the PI scores vary based on changes in two key metrics holding all else constant: (1) cost, which can take on values of High, Medium, and Low; and (2) Endangered Species Act (ESA) listing, which takes on values of Listed, Species of Concern, and Not Listed.

We analyze the impact of key metrics on PI scores for each species separately, using typical values for the *X*_*ij*_’s held constant, ascertained from our review of the WDFW manual (WDFW 2020). Detail on the metric levels is provided in the Appendix.

## Results

### Systematic Review

We identified 10 entities using culvert prioritization indexes: the City of Bellingham (Bellingham), Whatcom County (Whatcom), Skagit River System Corporative (Skagit), Snohomish County (Snohomish), King County (King), Thurston County (Thurston), Chehalis Basin (Chehalis), and Cold Water Connection Campaign (CWCC), and the Washington Department of Fish and Wildlife (WDFW) (Figure 1).

Among prioritizing entities, there was a high level of variation in terms of goals, geographic and jurisdictional scope, and metrics included the PI scores (Table S1). Each PI entity considered different culvert owners, some choosing to only score culverts owned by the PI entity, and some choosing to score all culverts regardless of ownership. In our sample, the number of scored barriers ranged from 32 to 4,259, the geographic extent ranged from 727 to 71,300 mi^2^, and the number of barrier owners considered ranged from 1 to 8. Four of these entities used the same non-linear formula, developed by the WDFW, for prioritization and five entities developed their own linear PI equation. The remainder of our results focuses on the six unique PI methods developed.

### All PIs: Distribution Results

The Shapiro-Wilk tests for normality identified Bellingham as the only normal (p <0.19) and negatively skewed distribution (Figure 2). The remainder of the PIs had a right skew, and WDFW had the greatest skew. Thurston county had the largest inter-quartile range (IQR) (32.6), and Chehalis had the smallest (9.06). A visual comparison shows that WDFW, Chehalis, and CWCC have evenly tapered tails, whereas Thurston and King have longer and more irregularly shaped tails.

**Figure 2.**
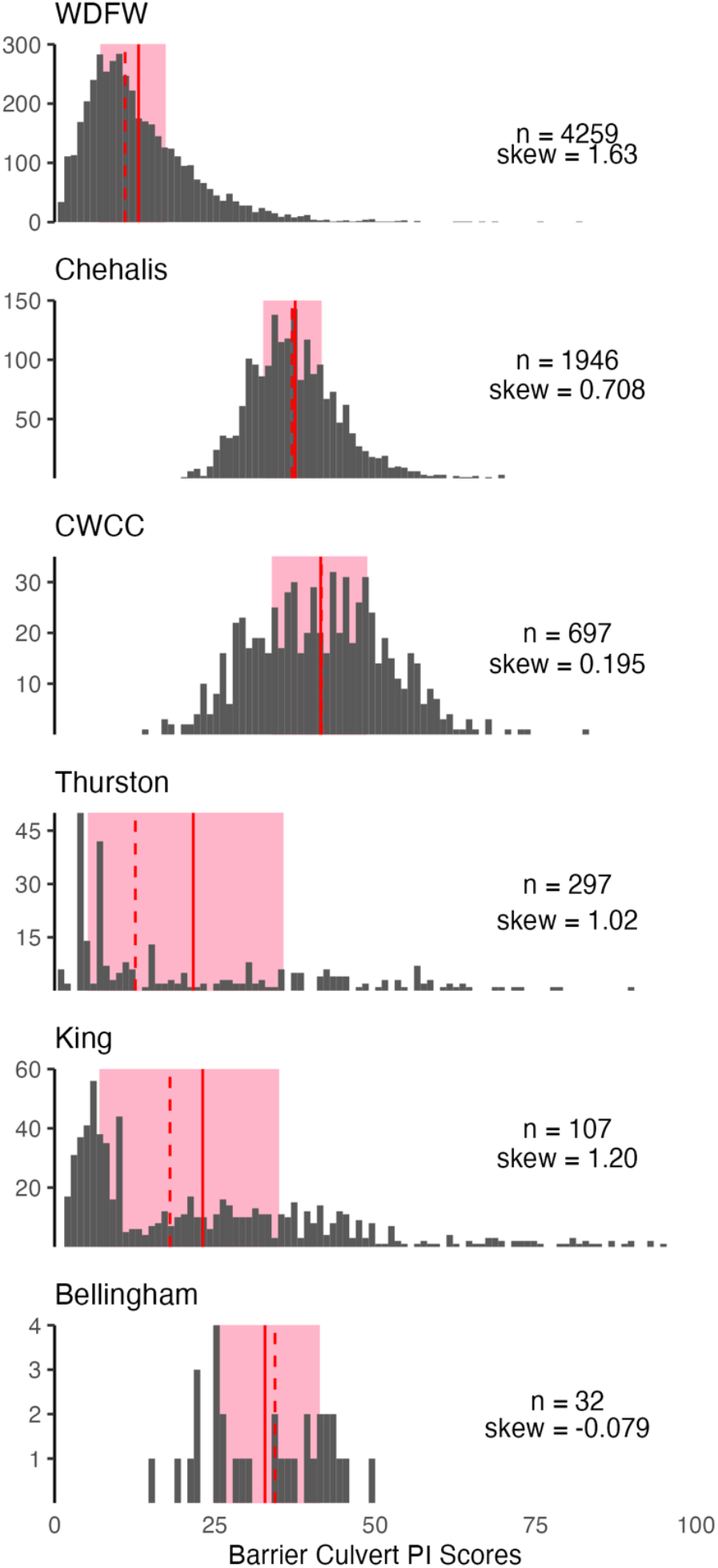
Histograms of the prioritization index (PI) scores for barrier culverts. The mean and median are displayed as a solid and dashed vertical line respectively. The IQR is shown as a shaded region. The histograms are ordered by sample size (n). All PIs were normalized to 100 as the maximum PI score. The scale of the y axis is determined by the maximum individual score frequency for each entity.

### All PIs: Shared Metric Analysis

First, we analyzed all metric definitions to determine whether the metrics used in the PI formulas were unique or shared with other entities. Overall, the number of unique metrics is greater than the number of shared metrics at 19 to 13 respectively. Next, using a qualitative metric comparison matrix, we plotted the percent number of shared metrics of each PI (Figure 3, A). We found that Thurston was the most unique PI, with the fewest shared metrics. Chehalis and WDFW were the least unique as all of their metrics were shared by at least one other PI studied.

**Figure 3.**
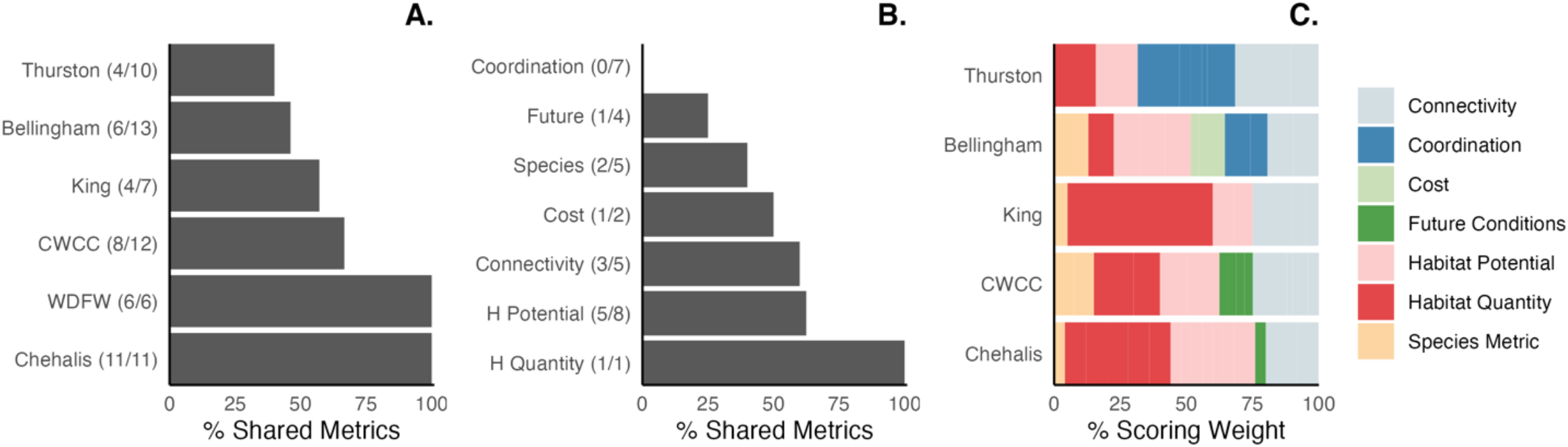
Bar graphs comparing the number of shared metrics and the scoring weight shown as a percentage. All PIs are included in Panel A and B but only the linear PIs are included in Panel C. Panel A shows the percent of shared metrics for each PI. Besides the PI entity name, in parenthesis, is the number of shared metrics/number of total metrics. Panel B shows the percent of shared metrics within each category of metric for all PIs studied. Besides the name of the category, in parenthesis, is the number of shared metrics/number of total metrics. Panel C shows the percent scoring weight of metric categories for each linear PI.

Finally, we pooled the PIs and analyzed the percent number of shared metrics within each category (Figure 3, B). We found that coordination was the most unique among categories of metrics and habitat quantity was the least. Table 2 provides further detailed information about the metrics coded as “shared” or “unique.” The most highly shared metrics were upstream habitat gain (100%), barrier passability (83%), and downstream barriers (83%).

**Table 2.** Summary of metric qualitative comparison matrix.

|  |  |
| --- | --- |
| **Metrics shared by 6 PIs:** | Upstream habitat gain |
| **Metrics shared by 5 PIs:** | Barrier passability, downstream barriers |
| **Metrics shared by 4 PIs:** | Species specific presence at barrier |
| **Metrics shared by 3 PIs:** | Upstream barriers, species IP value |
| **Metrics shared by 2 PIs:** | ESA listing, vegetation, road density, current stream temperatures, water quality 303d listing, road size as a proxy for cost, future stream temperature |
| **Unique metrics:** | Barrier clusters, barrier density, number of species with upstream habitat, mobility, juvenile benefits, other restoration projects, prioritized watershed, expert opinion on habitat quality, funding availability, other construction, community support, barrier condition, average traffic, traffic detour, maintenance, expert opinion on community impacts, future winter max flows, future winter bankfull flows, future summer flows |

### Linear PIs: Category Weights

Five of the PIs used a linear equation, with each metric as a weighted input. The similarity between linear equations across entities allowed for direct comparison of the scoring weight of metric categories (Figure 3, C). A visual comparison shows that PIs assign weights differently. For example, King assigned a relatively high weight to habitat quantity, whereas Bellingham assigned a relatively low weight to this category. Additionally, some entities included many categories, such as Bellingham, which results in overall reduced weights for individual categories.

### Non-Linear PIs: Weights

#### Scenario 1: Varying Costs

WDFW uses road class size as a proxy for cost, with larger roads considered higher cost, and smaller roads considered low cost. We model the change in the WDFW PI score, used by four entities, in relation to changes in barrier restoration cost using the median PI score (13) as a point of reference for comparison (Figure 4). We found that the scores exhibited a fixed ratio, in terms of upstream lineal habitat gain (ULHG) needed to achieve an equivalent PI score across cost scenarios (see the Appendix for a proof). All else equal, a low-cost culvert would need a third of the ULHG as a high-cost culvert to achieve the same PI score. Likewise, a medium cost culvert would need half the ULHG as a high-cost culvert to achieve the same PI score.

**Figure 4.**
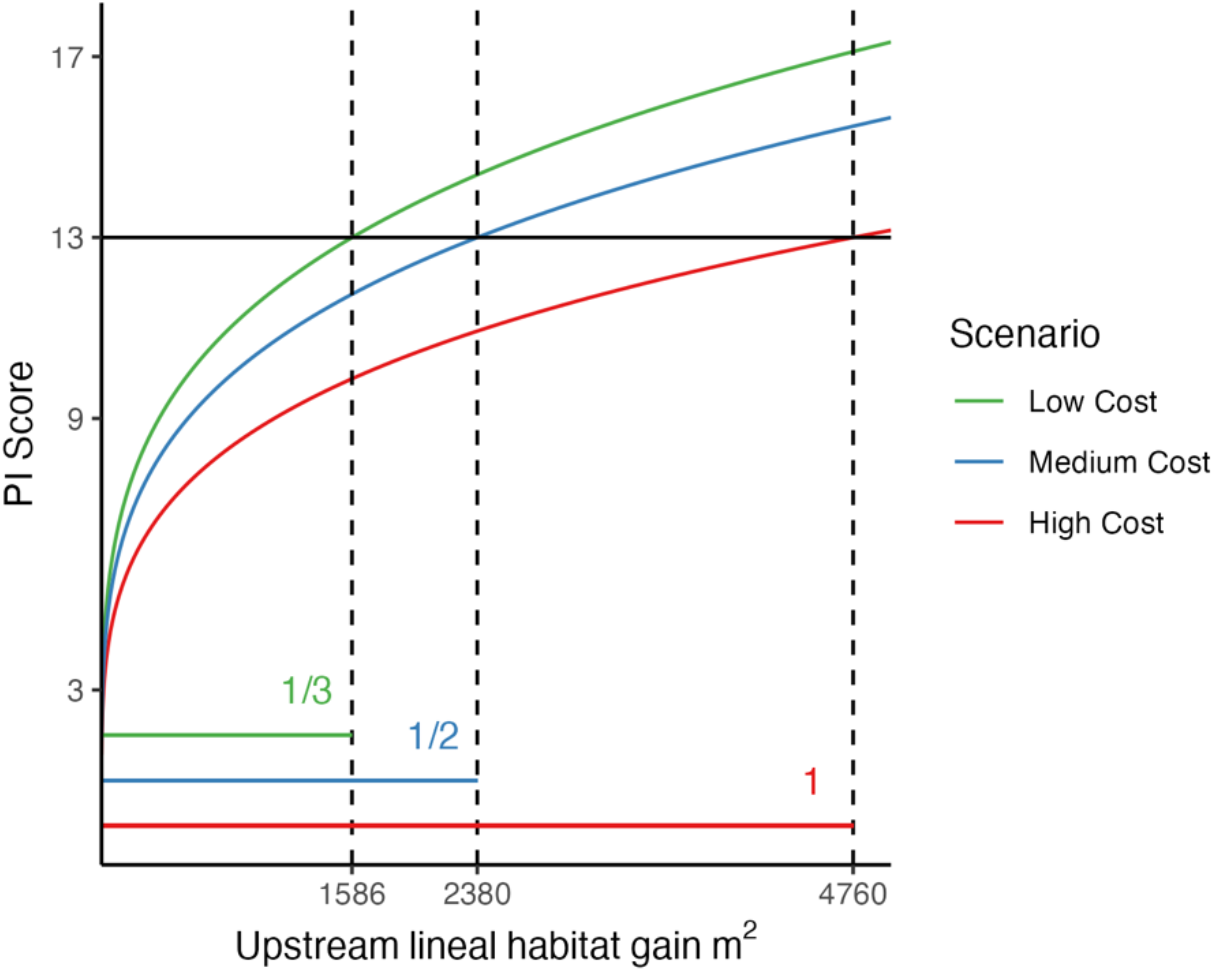
The impact of habitat gain for culverts of different cost categories on PI scores calculated with the WDFW method. The solid horizontal line shows the WDFW median PI score. Dashed lines show where the different cost scenarios intersect with the median PI score.

We further investigated the WDFW’s assumption on costs to see if the road class cost categories, i.e., a score of 1-3 for low, high, and medium cost projects respectively, fit data on restoration costs. Specifically, we examine cost ratios for medium- and low-cost barrier restoration projects as reported by the Pacific Northwest Salmon Habitat Restoration Project Tracking Database (PNSHP) data (Van Deynze et al. 2022, Pacific Northwest salmon habitat project database 2021). The PNSHP ratio of median costs for barrier correction projects on the middle-cost and lower-cost roads was 1.46, closely matching the ratio of 1.5 used by the WDFW (Table S3).

#### Scenario 2: Varying Species ESA Status

The WDFW formula accounts for ESA listed species through a metric that takes on three values (ESA Listed = 3, Species of Concern = 2, and Non-listed = 1). Thus, the ESA metric also generates a constant ratio of habitat required to equalize scores across ESA status (1/3 : 1/2 : 1) or (2 : 3 : 6), as seen in the previous cost scenario.

While the constant ratio property holds for ESA status, the X_ij_ metrics are species specific, with species that have a higher production potential (P) such as sockeye salmon (P = 3) having higher values for metrics in the habitat quality category than species with a low P such as bull trout/dolly varden (P = 0.0007). Therefore, the impact of ESA listing while holding habitat constant varies across species (Figure 5). Specifically, we find sockeye and chum salmon realize the largest PI score gains from a listing status of Endangered/Threatened (Figure 5). Production potential is also high for pink salmon, although there are no ESA-listed pink salmon populations in the Case Area (Figure 5).

**Figure 5.**
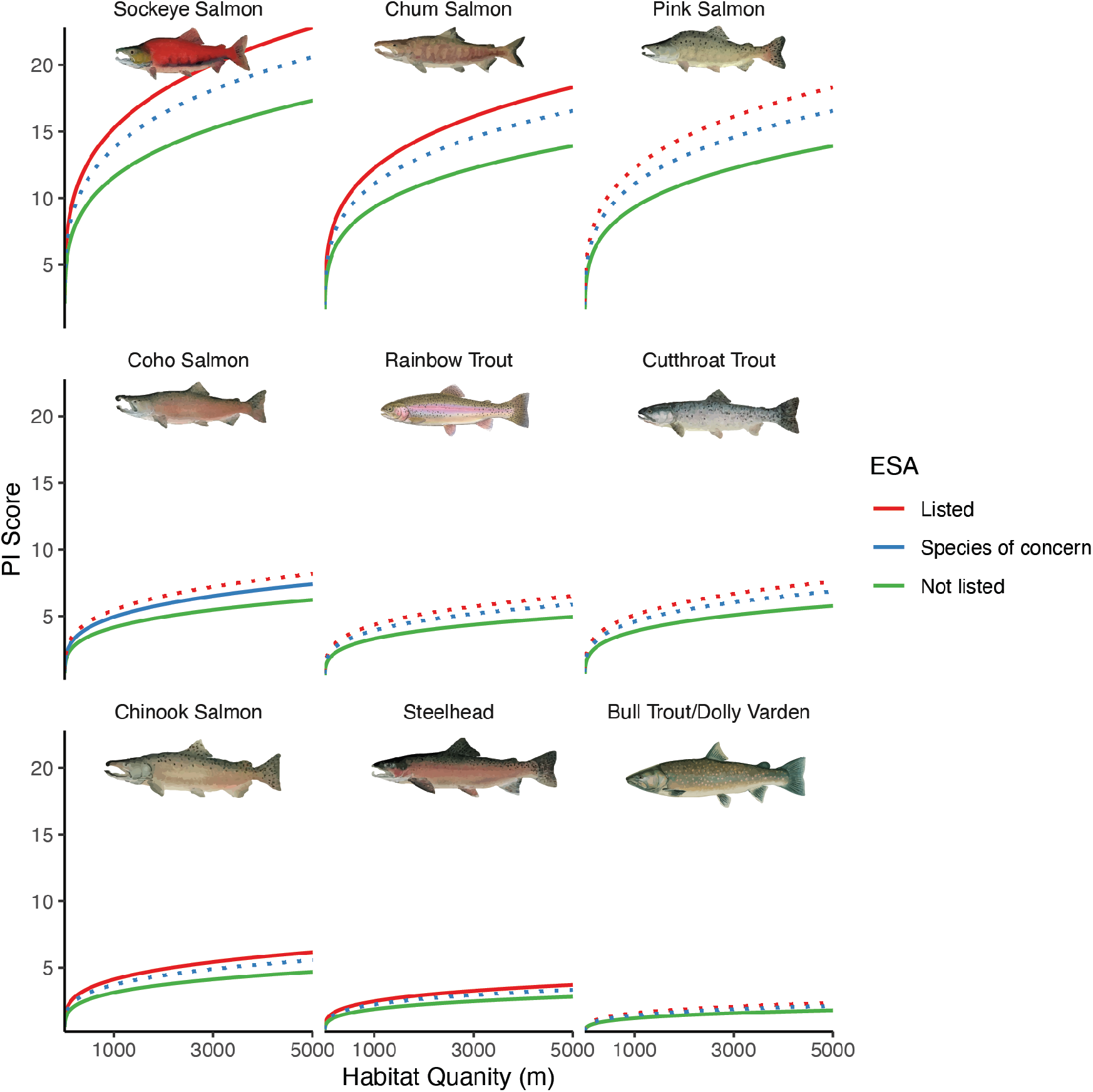
The impact of habitat gain for culverts of different ESA listing categories on PI scores calculated with the WDFW method. Species with ESA listed runs within the case area were modeled separately for ESA status with a solid line. The dashed line indicates possible scores for ESA listings that do not exist within the case area. Species were organized from the highest PI to lowest to demonstrate overall prioritization weight by species.

## Discussion and Conclusion

The need to replace barrier culverts to increase iconic salmon populations and uphold tribal treaty rights, is gaining traction across Western Washington, even for those entities not bound to do so by the federal injunction. As a result, in the last decade multiple prioritization frameworks have emerged in the region.

Here we conduct a comparative analysis of 6 prioritization frameworks adopted by 10 entities in Western Washington, together scoring an estimated 9,143 barrier culverts for prioritization. Our results increase the transparency of barrier culvert restoration priorities in the Case Area and demonstrate how attributes of a high-priority barrier differ across the entities.

We found that all entities in the Washington Case Area, have adopted PI scores to decide which projects to fund in the near term, e.g., in contrast to optimization methods or relying on expert opinion alone. Additionally, our study identified 5/10 entities using a common non-linear PI equation, developed by WDFW, and 5/10 entities that have developed a customized linear PI equation. Thus, there is variability in both the functional form of the PI formulas used as well as inputs to the function including variables and weights on those variables.

Government reports suggest that customization is driven by the feasibility for data collection, the number of culverts within an entity, presence or absence of anadromous species, and the desire to incorporate feedback from local experts, stakeholders, and tribal co-managers. Such widespread customization means that, even within a region, identical barrier culverts may be prioritized very differently, depending on barrier ownership.

There do exist some similarities between the PIs in the Case Area in terms of factors included. For example, Habitat Potential and Habitat Quantity are included in all of the PI formulas. Additionally, Connectivity is included in all 5 of the linear PI formulas, though the specific metrics used to measure these factors often varied.

Species Metric is included in 4/5 of the linear PI formulas and in the non-linear WDFW framework. In linear equations, Species Metrics were generally given low weight, while in the non-linear PI, the importance of ESA listing status (a type of Species Metric) depends on the productivity of the species, with sockeye and chum receiving the largest increase in PI from an ESA listing.

Surprisingly, cost is not included in 5/6 of the PI frameworks. Although, the WDFW framework is used by 5 entities and requires that high-cost barriers generate three times the amount of low cost barriers, a ratio consistent with estimated costs in the literature. In contrast, in Bellingham, the only entity with a linear PI framework including Cost, the scoring weight on Cost is 12.9%. This demonstrates that Bellingham’s cost metric has a lower influence on the overall score compared to the WDFW framework. Bellingham is also the only PI to factor in funding availability to their score.

Despite the fact that PI scores share both commonly included and excluded factors, the factors are typically represented by unique metric inputs, with the number of unique metrics employed outnumbering the number of shared metrics. Differences in metric inputs further demonstrates the ways in which barrier-owning entities customize their prioritization processes. For example, coordination was the most unique category, which included metrics for a range of collaborative examples such as educational opportunities, stakeholder engagement, or construction feasibility.

While there are clear benefits from customization including the ability of local entities to define their own priorities, utilize all of the data and information they have access to and determine to be valuable to the decision-making process, customization does introduce challenges in terms of transparency, coordination across entities (e.g. none of our PI scores give bonus points to barriers hydrologically connected to planned projects of other entities), and comparing candidate projects across entities as the range of scores vary across entities. Cross-entity comparisons are particularly important for external funding agencies seeking to allocate funds across the region to the projects with the greatest benefits.

It is important to note, however, written reports and personal correspondence has revealed that it is rare for entities to use PIs as the only deciding factor in barrier removal, and in reality, PIs are typically combined with other sources of information in a larger prioritization process. To our knowledge, these processes are undocumented, and we are unable to systematically assess what factors are included in the prioritization process outside of the PI scores. Thus, it is unclear whether these unobserved factors drive increased similarity in PI frameworks across the Case Area or increased divergence. For example, if cost information is as a part of the prioritization process for those 4/6 entities that do not include cost in their PI scores, differences may not be as great as they seem from an evaluation of the PI scores alone. On the other hand, the ability of the prioritization process to include local knowledge and local preferences, may exacerbate differences. Future research on the larger prioritization process is needed to more completely understand how entities are deciding which barrier culverts to fund in the near term with limited budgets.

## Acknowledgements

This project was funded by State funds from the University of Washington School of Marine and Environmental Affairs. We would also like to thank Daniel Howe of Snohomish County, Marcus Storvick and Trevin Taylor of Thurston County, and John Thompson of Whatcom County for providing us with information and/or data on the PI formula and how it fits in to culvert prioritization in their counties. We also thank participants at our virtual workshop, held in June of 2022, funded by the Washington Sea Grant (grant #NA22OAR4170103), and entitled “Fish Passage Planning in Washington: A Decision Support Tool”. All mistakes are our own.

