## Supplementary Tables for "Western Washington culvert prioritization indexes: A cross jurisdictional comparative analysis of management strategies for salmon recovery"

### Supplementary Figures

**Table S1.** General information extracted from search and screen process.

| PI Developer | Ownership Extent | Geographic Extent | Developed Unique Prioritization Method | Number of Barrier Culverts with PI Scores | Interactive Tool |
|---|---|---|---|---|---|
| Chehalis Basin - WDFW[1] | State, Tribal, Private, County, Federal, City | WRIA 22 & 23: 2766 $mi^2$ | Yes | 1946 | Yes |
| City of Bellingham[2] | City | City of Bellingham: 30.51 $mi^2$ | Yes | 32 | No |
| Cold Water Connection Campaign / Coast Salmon Partnership[3] | State, Tribal, Private | WRIA 20 & 21: 2850 $mi^2$ | Yes | 697 | Yes |
| King County[4] | County | King County: 2,307 $mi^2$ | Yes | 107 | Yes |
| Skagit River System Cooperative (SRSC)[5,6] | State, County, Federal, City, Other and Private | Skagit River Basin: ~2,656 $mi^2$ | No | 443 | No |
| Snohomish County | County | Snohomish County: 2,196 $mi^2$ | No | 536 | No |

DRAFT

[1] Dwight, C. (n.d.). *Chehalis Fish Passage Barrier Prioritization* [Presentation]. https://www.chehalisbasinstrategy.com/wp-content/uploads/2020/01/7_Chehalis-Fish-Passage-Barrier-Prioritization-Model.pdf

[2] Burns, A. (December 15, 2019). *2019 City of Bellingham Fish Barrier Prioritization Update.* Public Works Department, City of Bellingham. https://cob.org/wp-content/uploads/2019-fish-barrier-prioritization.pdf

[3] Zimmerman, M. (December 31, 2020). *Assessment and Restoration Design of County Road Fish Barriers on the Western Olympic Peninsula.* Coast Salmon Partnership.

[4] King County. (March 2021). *Final Report Regarding Remedies to Existing Fish Passage Barriers for King County.* https://your.kingcounty.gov/dnrp/library/water-and-land/habitat-restoration/fish-passage-restoration/remedies-to-existing-fish-passage-barriers-report.pdf

[5] Mickelson, E., Smith, D., Hinton, S. (June 7, 2020). *Skagit Basin Barrier Culvert Analysis: Public and Private Stream Crossings. http://skagitcoop.org/wp-content/uploads/Skagit-Basin-Barrier-Culvert-Analysis-Report-and-Appendices.pdf*

[6] The Watershed Company. (January 2014). *Skagit County High Priority Culvert Replacements for Fish Passage. https://www.skagitcounty.net/PublicWorksNaturalResourcesManagement/Documents/High%20Priority%20Culvert%20Replacements%20for%20Fish%20Passage.pdf*

| | | | | | |
|---|---|---|---|---|---|
| Thurston County[7] | County | Thurston County: 727 mi² | Yes | 279 | Yes |
| Washington Department of Fish and Wildlife (WDFW)[8] | State, Tribal, Private, County, Federal, City, Port, Drainage District, Irrigation District | State wide: 71,300 mi² | Yes | 4259 | Yes |
| Washington State Department of Transportation (WSDOT)[9] | State | State wide: 71,300 mi² | No | 366 | No |
| Whatcom County[10] | State, County, City, Private | Whatcom County: 2,503 mi² | No | 478 | No |

DRAFT

7 Thurston County Public Works. (n.d.). *Fish Passage Enhancement.* *https://www.co.thurston.wa.us/publicworks/Projects/63000/Fish%20Passage%20Enhancement%20Program.pdf*

8 Barrett, D., Zweifel, J. (2019). *Fish Passage Inventory, Assessment, and Prioritization Manual.* Washington Department of Fish and Wildlife. *https://wdfw.wa.gov/publications/02061*

9 Kanzler, S., Romero, D., Prosser, K., Schmidt, T., Hershfield, M. (June 30, 2021). *WSDOT Fish Passage Performance Report.* WSDOT. https://wsdot.wa.gov/sites/default/files/2021-10/Env-StrRest-FishPassageAnnualReport.pdf

10 Whatcom County. (January 2006). *Whatcom County Fish Passage Barrier Inventory Final Report.* IAC Project Number: 01-1258 N. https://salmonwria1.org/sites/default/files/2019-09/A%20Final%20Report_culvert%20inventory.pdf

**Table S2:** Detailed inputs to PI equations and metric categorization.

| PI Developer | Method | Species Metrics | Habitat Quantity Metrics | Habitat quality Metrics |
|---|---|---|---|---|
| **King County** | Linear equation | **1. Species benefits (0, 5)** for Chinook or Sammamish Kokanee. | **1. Habitat Gain (0-55)** determined by a 200m upstream stream survey for Coho rearing intrinsic potential (IP). The 200m is broken down into segments that fall into Coho IP reach scores. The binned IP score is multiplied by the number of stream segments, summed for the whole 200m and normalized to 55 total points. | **1. Percentage forested (0, 1, 3, 5, 8)**<br>**2. Percentage impervious (0, 1, 3, 5, 7)** |
| **Cold Water Connection Campaign / Coast Salmon Partnership** | Linear equation | **1. Documented species (0:6)** measures the presence of priority species. +1 for each fish present including Coho, Steelhead, Chinook, Sockeye, Bull trout, Resident<br>**2. Species Habitat (0:6)** measures the presence of upstream habitat for priority species.<br>+1 for each fish present including Coho, Steelhead, Chinook, Sockeye, Bull trout, Resident | **1. Anadromous Habitat Gain (0:5)** the linear upstream habitat in miles to next 0% passable barrier or end of species distribution whichever is shorter. This is calculated independently for species of interest, and the species with the highest value is used for choosing bin number.<br>**2. Resident Habitat Gain (0:5)** is the same metric applied to resident species of interest. | **1. Intrinsic Potential (1:5)** score from upstream habitat segments stopping at 0% passable barrier or end of fish distribution. IP is calculated for Coho, Chinook, and Steelhead.<br>**2. Current Stream Temperatures (1:5)** are based on optimal temperature ranges for fish species of interest. |
| **Chehalis Basin (WDFW)** | Linear equation | **1. Number of target species (1:5)** benefitting from the project. Species include Chinook, steelhead, coho, searun cutthroat, and chum. | **1. Habitat Gained (0:10)** is scored separately for each of the five target species. This variable measures the amount of miles of upstream habitat gained. | **1. Road density (1:5)** measures $mi/mi^2$ of roads in proximity to barrier.<br>**2. Water quality (1, 5)** is a binary variable that considers if the site has a 303d listing upstream.<br>**3. Stream temperature (0:5)** uses optimal temperature range.<br>**4. Riparian vegetation (1:10)** is calculated based on canopy cover, average tree height (m) and average buffer width (m).<br>**5. Intrinsic potential (0:5)** is scored for each target species and summed along with the habitat gain metric. |
| **Washington Department of** | Quadratic root | A unique PI is calculated for each species present at culvert, these individual PI | **1. Habitat Gained (0-72,607)** | **1. Habitat Quality Metric (HQM) (0.33, 0.67, 1)** calculated from physical habitat surveys that |

DRAFT

| | | | | |
|---|---|---|---|---|
| **Fish and Wildlife (WDFW)** | (geometric mean) | scores are summed together to get the total PI score.<br>**1. The species mobility (1, 2)** includes if the species is anadromous or resident.<br>**2. The species condition (1:3)** incudes if the species/run is ESA listed, a species of concern, or not listed. | is measured in upstream lineal gain in m^2. This variable is continuous. | measure habitat type, substrate type, thermal cover, in stream cover and temperatures and other metrics. The HQM measures the availability of spawning and rearing habitat upstream from the barrier. Species that are spawning limited (chum, sockeye/kokanee, pink) will use the spawning HQM. Species that are rearing limited (Coho, Chinook , cutthroat, steelhead, resident trout, and bull trout/Dolly Varden) will use the rearing HQM.<br>**2. Intrinsic potential (0.0007, 0.0021, 0.016, 0.037, 0.04,, 0.05, 1.25, 3)** species specific fixed constant for the production capacity for that species per meter squared of habitat. |
| **City of Bellingham** | Linear equation | **1. Species condition (1:3)** considers the number of ESA listing of species present, >2 listed species, 1 listed species, no listed species.<br>**2. Juveniles metric (0, 1)** considers if there are anadromous juveniles present. | **1. Habitat Gained (0:3)** is a single variable measured in upstream lineal gain in m. | **1. Proximity to other restoration projects (0.5, 1, 1.5, 2, 2.5, 3)** measured in feet.<br>**2. Surface water benefit (0, 3)** Increased flood storage +1, expanded floodplain +1, incorporates 303d waterway +1<br>**3. Watershed prioritization (1, 2, 3)** Priority watershed region, Tier 1 sub-watershed, Tier 2 sub-watershed |
| **Fish Passage Enhancement Program (Thurston County)** | Linear equation | **1. Species metric (0, 1)** is multiplied into the total PI score. If there are salmon present then the PI score will be run. If there are no salmon present then the PI score will be zeroed out. | **1. Habitat Gained (0, 2, 5, 7, 10, 15)** is measured in lineal gain upstream in feet. | **1. Habitat quality (0, 7, 15)** is scored qualitatively as high, medium, low/other. |

DRAFT

| PI Developer | Connectivity Metrics | Coordination Metrics | Cost Metrics | Future Conditions Metrics |
|---|---|---|---|---|
| **King County** | **1. Downstream barriers (1, 4, 7, 10, 15)** considers the number and passability of downstream barriers.<br>**2. Barrier clusters (0, 5)** considers if the culvert is in proximity to a group of >3 barriers and >15 barriers/mi and 704 ft maximum distance, or barrier with pipe >1,000ft.<br>**3. Barrier density (0, 3, 5)** is determined by identifying furthest downstream barrier, counting the number of barriers in subbasin upstream, totaling the length of IP watercourse in sub basin, dividing the # of barriers by length of IP. | N/A | N/A | N/A |
| **Cold Water Connection Campaign / Coast Salmon Partnership** | **1. Barrier passability (1:5)** is the percent passability of fish through the barrier.<br>**2. Number of downstream barriers (0:5)**<br>**3. Passability of downstream barriers (0, 2, 4, 5)**<br>**4. Number of upstream barriers (0:5)**<br>**5. Passability of upstream barriers (0, 2, 4, 5)** | N/A | N/A | **1. Future winter max flows (1:5)** looks at percent change from historic to 2080 prediction.<br>**2. Future winter bankfull flows (1:5)** considers percent change from historic to 2080 prediction.<br>**3. Future summer flows (1:5)** looks at the percent change from historic to 2080 low flow prediction.<br>**4. Future stream temperature (1:5)** scores based on optimal temperature ranges. |
| **Chehalis Basin (WDFW)** | **1. Barrier passability (5, 10, 15)** is the general percent passability of fish through the barrier.<br>**2. Downstream barriers (0, 3, 4, 5)** uses both quantity (0, 1, 2, >2) and percent passability.<br>**3. Upstream barriers (0, 3, 4, 5)** uses both quantity (0, 1, 2, >2) and percent passability. | N/A | N/A | **1. Future stream temperature (0:5)** is scored based on optimal temperature ranges. |

DRAFT

| | | | | |
|---|---|---|---|---|
| **Washington Department of Fish and Wildlife (WDFW)** | **1. Barrier passability (0.33, 0.67, 1)** is the general percent passability of fish through the barrier. | N/A | **1. Cost modifier (1:3)** is applied based on the size of the road. Private/single lane road (low cost), City/county road or state 2 lane road, state highway >2 lanes (high cost) | N/A |
| **City of Bellingham** | **1. Barrier passability (0.5, 2, 3)** is the general percent passability of fish through the barrier.<br>**2. Proximity with other planned/completed barriers removals (0.5, 1, 1.5, 2, 2.5, 3)** upstream and downstream is measured in feet. | **1. Construction coordination (0:3)** considers if the culvert is at the same location as a similar construction project by 2025.<br>**2. Community support (0:2)** considers education opportunities and willingness of stakeholders.<br>Educational opportunities +1, willing stakeholders +1 | **1. Funding opportunities (0, 1)** outside the Fish Barrier Removal Board +1.<br>**2. Cost estimations (0:3)** are based on the width of the proposed structure. | N/A |
| **Thurston County** | Connectivity includes two variables. Barrier passability is the general percent passability of fish through the barrier.<br>0% = 20, 33% = 15, 67% = 10, Unk/Other = 0<br>Downstream barriers is a binary presence/absence variable.<br>No = 10, Yes = 5, Unk = 0 | **1. Barrier condition (0, 5, 7, 10, 15)** is scored by the barriers structural defects. Barriers with high structural defects are prioritized.<br>**2. Average Daily Traffic (0, 4)** is scored by the busyness of the road, giving higher scores to busier roads. The reason is because it provides a greater benefit to the local community versus a rural gravel road with lower traffic volumes.<br>**3. Traffic detour (0, 4)** is a binary variable (yes/no).<br>**4. Maintenance score (0, 1, 2)** prioritizes culverts that require frequent maintenance.<br>**5. Expert score (0, 5, 10)** is a qualitative priority score designated by a professional judge. Once the review has been completed for a potential project the Thurston Staff has two Biologist assigned with over 40 years of experience to make the final determination. | N/A | N/A |

DRAFT

**Table S3.** Cost summary statistics by road category. Road class 3 matches WDFW's proxy for low cost, road class 2 is WDFW medium cost proxy, and road class 1 is WDFW high cost proxy. Note that there are no barriers in the Case Area in the PNSHP sample that fall within the WDFW Class 3 definition.

| Road Class | n | Mean Cost | Median Cost | Standard Deviation Cost |
|---|---|---|---|---|
| 3 | 46 | 129,305 | 86,816 | 117,440 |
| 2 | 83 | 188,974 | 126,333 | 177,295 |
| 1 | 0 | N/A | N/A | N/A |

**Proof of WDFW cost and species condition ratios.** Both cost and species condition (ESA listing status) can take the value of 1, 2, or 3. To demonstrate the constant ratio we hold the PI score constant at a value of P and define $H_i$ as the upstream habitat gain needed to achieve a PI score of P when cost (or species condition) takes on a value of i $\in \{1, 2, 3\}$ when all other variables are equivalent and set to $\theta$.

$$P = \sqrt[4]{\theta H_1} \implies P^4 = \theta H_1,$$

$$P = \sqrt[4]{\theta 2 H_2} \implies P^4 = \theta 2 H_2\,,$$

$$P = \sqrt[4]{\theta 3 H_3} \implies P^4 = \theta 3 H_3.$$

This implies:

$$\frac{H_1}{2} = H_2\,,$$

and

$$\frac{H_1}{3} = H_3$$

DRAFT
